# Selective Activation of Macrophage Innate Signaling Pathways and Inflammatory Responses to *Orientia tsutsugamushi* Karp and Gilliam Strains

**DOI:** 10.64898/2026.04.24.720537

**Authors:** Dario Villacreses, Casey Gonzales, Yuanyi Zhang, Yuejin Liang, Lynn Soong

**Author notes:** Equal contributions and address correspondence to: Lynn Soong, Yuejin Liang.

## Abstract

*Orientia tsutsugamushi* (*Ot*) is an obligatory intracellular bacterium that can cause scrub typhus, an emerging but severely neglected disease with high mortality rates. *Ot* Karp and Gilliam strains account for most reported cases in Southeast Asia. Our group has reported that Karp-infected outbred and inbred mice exhibit more severe disease outcomes than their Gilliam-infected counterparts, likely due to their excessive inflammation and tissue injury. Macrophages (MΦs) serve as the main target cells for *Ot* replication and host defense against the infection; they also are key players in immune modulation and cytokine/chemokine production. However, it remains unclear as to how MΦ-*Ot* interactions impact disease outcomes. In this study, we focused on RNAseq from C57BL/6 mouse-derived MΦs to reveal *Ot* infection*-* and strain-related immune signatures. While *Ot* infection modulated several common canonical pathways/genes, we identified unique pathway/gene signatures that were highly selective for Karp or Gilliam strain, respectively. Karp infection uniquely upregulated proinflammatory signaling and pattern recognition receptors, including C-type lectin receptors (CLRs), including Mincle/*Clec4e* and Dectin-2/*Clec4n*. In contrast, Gilliam infection enhanced MΦ proliferation and DNA replication, and Gilliam strain grew better in M0-like MΦs than Karp strain. In IFN𝛾-primed (M1-like) MΦs, however, Karp exhibited significantly greater resistance against host killing than Gilliam, suggesting its superior ability to evade host immune responses. Overall, Karp strain preferentially upregulated CLRs and activated type 1-skewed inflammatory responses, but it is also relatively resistant to IFN𝛾-mediated killing. This study provides new insights into potential mechanisms underlying *Ot* strain-associated immune responses and disease outcomes.

**Author Summary:** *Orientia tsutsugamushi* (*Ot*), the causative agent of scrub typhus, is an understudied, life-threatening pathogen endemic to Southeast Asia. Among its various pathogenic strains, Karp and Gilliam are clinically dominant and exhibit distinct disease severity. Building on previous reports that Karp-infected mice display more severe disease than those infected with Gilliam, we explored how strain-specific interactions with macrophages (MΦs), a central cell type in both *Ot* replication and host immunity, may contribute to these outcomes. Using RNAseq of primary MΦs from C57BL/6 mice, we characterized transcriptional profiles of both strains. Karp infection was uniquely associated with increased proinflammatory signaling and the selective induction of bacterial-sensing molecules. In contrast, Gilliam infection promoted host cell proliferation and supported more efficient bacterial replication in naïve MΦs. Notably, Karp demonstrated increased resistance to host killing in stimulated (proinflammatory) MΦs compared to Gilliam. These findings suggest that the Karp strain can drive strong inflammatory responses but is adept at evading IFN𝛾-mediated immune defenses. Our study uncovers key immunological distinctions between *Ot* strains and offers novel insights into the mechanisms of strain-specific pathogenesis in scrub typhus, with potential to uncover biomarkers for disease severity.

## Introduction

*Orientia tsutsugamushi (Ot)* is a unique but poorly studied intracellular bacterium species, causing scrub typhus in humans. Unlike closely related bacteria in the *Rickettsia* genus, *Ot* exhibits a distinct outer membrane architecture (with its outer leaflet thicker than the inner leaflet), expresses low levels of peptidoglycan-like structure*s*, and totally lacks lipopolysaccharides, which are key ligands for TLR4/TLR2-mediated immune recognition (1). These structural differences are known to influence host-pathogen interactions and may contribute to variations in immune recognition (2–4). However, our understanding of *Ot*-induced cellular and molecular details and their impact on disease outcomes remains obscure, partially due to the lack of *Ot*-based genetics tools (5).

*Ot* has historically been endemic to Southeast Asia; yet growing evidence has revealed the detection of its DNA and/or antibodies in the Middle East, Europe, Africa, and South and North America (6, 7). While most infected cases are self-limiting, scrub typhus is a life-threatening disease, which can lead to ∼30% mortality rates in certain countries/regions due to misdiagnosis or mistreatment (8). Lethal outcomes are associated with phagocyte and endothelial cell infection in multiple organs (lungs, spleen, liver, brain, etc.), vascular dysfunction (9, 10), and multi-organ dysfunction syndrome (11). At present, over 40 *Ot* genotypes and 20 serotypes are known to be pathogenic in humans (12, 13). In Southeast Asia, Karp and Gilliam strains (or serotypes) account for approximately 50% and 25% of cases, respectively (13, 14). Molecular analysis of these strains has shown differences within the variable region of the TSA56 gene, which may contribute to antigenic variation. Furthermore, differences in virulence and pathogenicity have also been reported among strains, highlighting the complexity of *Ot* infections (15–17). Recent human studies suggested that Karp infection has higher virulence than Gilliam infection, likely due to differential immune responses (18). Non-human primate studies suggested that skin-inoculated Gilliam resulted in more progressive skin lesions than Karp group(19, 20). Yet detailed immunological studies in humans or in non-human primates, either at the cellular or molecular level, remain very limited.

The recently developed murine scrub typhus models and comparative studies have revealed valuable insights into host immune responses and disease outcomes (17, 21–23). For example, in comparative studies with two *Ot* strains (Karp vs. Gilliam) and two mouse models (outbred CD-1 vs. inbred C57BL/6 mice), our group found that Karp infection induced stronger host susceptibility and inflammatory immune responses than Gilliam infection, regardless of mouse models and inoculation doses (17, 23). Notably, while Karp-infected CD-1 mice (100%) succumbed to infection by day 15, only 50% of Gilliam-infected mice died, with the rest of the mice starting body weight recovery at day 15 (23). Even though we identified susceptibility differences between murine models, Karp strain consistently presented higher pathogenesis compared to Gilliam (17). These findings underscore strain-dependent differences in pathogenesis and highlight the need to define the molecular and cellular determinants that govern these divergent outcomes. Elucidating these mechanisms will provide a framework for understanding how immune regulation is balanced between host protection and pathology, thus defining *Ot* strain-related cellular pathways and mechanisms underlying the diverse immune responses and disease outcomes.

Macrophages (MΦs) are one of the main targets and effector cells during *Ot* infection and play complex roles. MΦs can act as target cells during initial infection, facilitating pathogen dissemination to local lymph nodes, followed by systemic spread (24, 25). Recent reports have described how *Ot* may induce excessive neuroinflammation driven by microglia (brain MΦs), accompanied by blood-brain barrier dysregulation during severe stages of infection in mice (10). MΦs also play a central role in immunity by releasing immune mediators (cytokines, chemokines, etc.) that direct immune responses (26), but their dysregulation can also drive inflammation and tissue damage (27). Following the analysis of lung MΦ subsets in Karp-infected mice, we have revealed an M1-type polarization of MΦ in the lungs of severe scrub typhus (22). Furthermore, in vitro studies have further revealed potential roles of MΦ subsets in *Ot* replication, as M2-type MΦs failed to control bacterial growth compared to M1-type cells (22). A better understanding of how different *Ot* strains interact with MΦs is crucial to defining differential host immune responses and disease outcomes.

Pattern recognition receptors (PRRs) comprise a diverse group of germline-encoded proteins that enable the innate immune system to detect conserved molecular signatures associated with pathogens or cellular damage. Among the major PRR families, Toll-like receptors (TLRs), C-type lectin receptors (CLRs), and NOD-like receptors (NLRs) play central and complementary roles in immune surveillance and host defense (28). TLRs are membrane-bound receptors located on the cell surface or within endosomal compartments, where they recognize a wide range of microbial components, including nucleic acids and lipoproteins, and initiate signaling cascades that promote inflammatory and antiviral responses. In contrast, CLRs are primarily involved in the recognition of carbohydrate structures in a calcium-dependent manner (29), contributing to pathogen uptake, antigen presentation, and modulation of immune responses, particularly in myeloid cells (30, 31). NLRs, which reside in the cytoplasm, detect intracellular perturbations and microbial motifs, often leading to inflammasome assembly and activation of proinflammatory cytokines such as IL-1β (4, 32). In the context of *Ot* infection, several PRRs like TLR2 and Mincle/*Clec4e* have been identified as immune sensor candidates (2, 21, 33). However, very little is known about the selective activation of these innate sensors by different *Ot* strains, nor their downstream pathways triggered by clinically prevalent *Ot* strains.

In this study, we compared transcriptomic profiles during Karp vs. Gilliam infection in vivo and in vitro to identify common pathways/genes shared across *Ot* infection and, more importantly, unique signatures specific to each strain infection. Consistent with our previous studies with Karp- vs. Gilliam-infected mouse tissues (17, 34), we found that Karp infection preferentially upregulated proinflammatory pathways/genes and host defense genes in the lungs and in MΦs. Notably, Gilliam infection primarily upregulated pathways/genes involved in cell cycle progression, DNA replication/repair, and cell proliferation. Moreover, we found a differential degree of resistance to IFNγ-mediated killing between Karp vs. Gillam strains. This study highlights distinct PRR gene expression profiles induced by different *Ot* strains, with a clear, distinct activation of CLRs, suggesting potential *Ot* strain-specific bacterial sensing patterns and regulatory pathways to manipulate immune responses and disease outcomes.

## Materials and Methods

### Mouse Infection and Tissue Collection

Female C57BL/6J (B6) mice were purchased from Jackson Lab Inc. Animals were infected at 8-12 weeks of age, following Institutional Animal Care and Use Committee-approved protocols (IACUC#2101001A) at the University of Texas Medical Branch (UTMB) in Galveston, TX. All infections were performed in UTMB ABSL3 facilities in the Galveston National Laboratory, and subsequent tissue processing and/or analysis were performed in BSL3 or BSL2 facilities, respectively. UTMB complies with the USDA Animal Welfare Act (Public Law 89-544), the Health Research Extension Act of 1985 (Public Law 99-158), the Public Health Service Policy on Humane Care and Use of Laboratory Animals, and the NAS Guide for the Care and Use of Laboratory Animals (ISBN-13). UTMB is registered as a Research Facility under the Animal Welfare Act and has current assurance with the Office of Laboratory Animal Welfare in compliance with NIH policy. *Ot* Karp and Gilliam stocks prepared from L929 cells were utilized for infection, as in our previous reports (17, 34). Briefly, B6 mice were intravenously inoculated with 5.6 × 10^4^ focus-forming units (FFU, in 200 μl) of Karp, Gilliam, or PBS (a mock control). Lung tissues (5/group) were collected at 4, 8, and 12 days post-infection (dpi) or from mock animals and inactivated for immediate or subsequent analyses. Disease severity was assessed using a standardized clinical scoring system ranging from 0 to 5, under an established animal sickness protocol (22). This scoring system incorporated multiple clinical parameters, including activity level, posture, fur condition, presence of bilateral conjunctivitis, and degree of weight loss, as outlined in our previous publication (17).

### Generation and Infection of Mouse Bone Marrow-Derived MΦs

Bone marrow cells were collected from the tibia and femur of mice and treated with red blood cell lysis buffer (Sigma Aldrich). MΦ were generated by incubating bone marrow cells at 37°C with 40 ng/ml M-CSF (Biolegend, San Diego, CA) in complete RPMI 1640 medium (Gibco) (22). Half of the culture medium was replenished on days 3 and 6, respectively; cells were collected on day 8. Viable cells were seeded into 24-well plates (5 x 10^5^ cells/well) or 6-well plates (5 x 10^5^ cells/well) for overnight incubation. To generate an inflammatory-like MΦ phenotype, cells were treated with 100 ng/ml IFN𝛾 (Biolegend) 24 h before infection. Bacteria were added at a multiplicity of infection (MOI) of 2 or 10, depending on experiments, with triplicate per condition. The same Karp or Gilliam stocks prepared from L929 cells were used in replicate experiments to ensure consistency.

### RNA-seq Assay and Data Analysis

Total RNA was extracted using RNeasy RNA Isolation kits (Qiagen). Samples (3 cultures/group) were sent to LC Sciences (Houston, TX) for RNA purity/quantity assessment using the Bioanalyzer 2100 and RNA 6000 Nano LabChip Kit (Agilent), followed by mRNA extraction, cDNA library construction, and sequencing with the Illumina Novaseq™ 6000. The Mus musculus genome build mm10 was used as a reference genome. Quality control analysis was done by using FastQC and RSeQC6. The company performed the initial differential expression analysis, as well as GO enrichment and KEGG enrichment analysis, as part of the service. Heatmaps were generated from differential expression analysis using GraphPad Prism.

Canonical pathway analysis was performed using QIAGEN Ingenuity Pathway Analysis (IPA), with methods and algorithms described by Krämer et al. (35). Genes with an |log2(FC)| > 1.25 and a *q*-value < 0.05 were included in analyses. Pathways significantly associated with the dataset were identified from the IPA canonical pathway library. Statistical significance was assessed using the Benjamini-Hochberg procedure, with pathways considered significant at a *q*-value < 0.05. Two separate scatter plots were generated: one for upregulated genes and one for downregulated genes, displaying the corresponding BH p-values for each pathway. Data analysis and visualization were conducted using R4.4.3 (35).

### Quantitative Reverse Transcriptase PCR (qRT-PCR)

To determine host gene expression, total RNA of MΦ was extracted via RNeasy mini kit (Qiagen), and cDNA was synthesized utilizing an iScript cDNA kit. qRT-PCR assays were performed using iTaq SYBR Green Supermix (Bio-Rad) on a CFX96 Touch Real-Time PCR Detection System (Bio-Rad). The assay included denaturing at 95ΦC for 3 min followed by 40 cycles of 10s at 95ΦC and 30s at 60ΦC. To check the specificity of amplification, melt curve analysis was performed. Transcript abundance was calculated utilizing the 2^-ΦCT^ method and normalized to *Gapdh*.

### Bacterial Burden Analysis (qPCR)

To determine bacterial loads, MΦ DNA samples were collected 3 dpi via a DNeasy kit (Qiagen) and used for qPCR, as described previously (21). Bacterial loads were normalized to the bacterial copy number per well. Data are expressed as a copy number of the 47-kDa gene per ng of DNA. The copy number of the 47-kDa protein gene was determined by using known concentrations of a control plasmid harboring a single-copy insert of the gene. A serial dilution of the control plasmid was then used to determine sample gene copy numbers.

### Statistical Analysis

Data were analyzed using GraphPad Prism software and presented as mean Φ SD. Differences between control and treatment groups were analyzed using one-way ANOVA with Šídák’s multiple comparisons test for Figure 1, Two-way ANOVA with Tukey’s multiple comparisons test, comparing groups only within the same time point for Figure 4, One-way ANOVA with Tukey’s multiple comparisons test for Figure 5B, and Two-way ANOVA with Tukey’s multiple comparisons test, comparing groups only within the same time point for Figure 5C. #*p* < 0.05, ##*p* < 0.01, ###*p* < 0.001, and ####*p* < 0.0001, respectively. Statistically significant values in comparison to *Ot*-infected samples are denoted as \**p* < 0.05, \*\**p* < 0.01, \*\*\**p* < 0.001, and \*\*\*\**p* < 0.0001.

**Figure 1.**
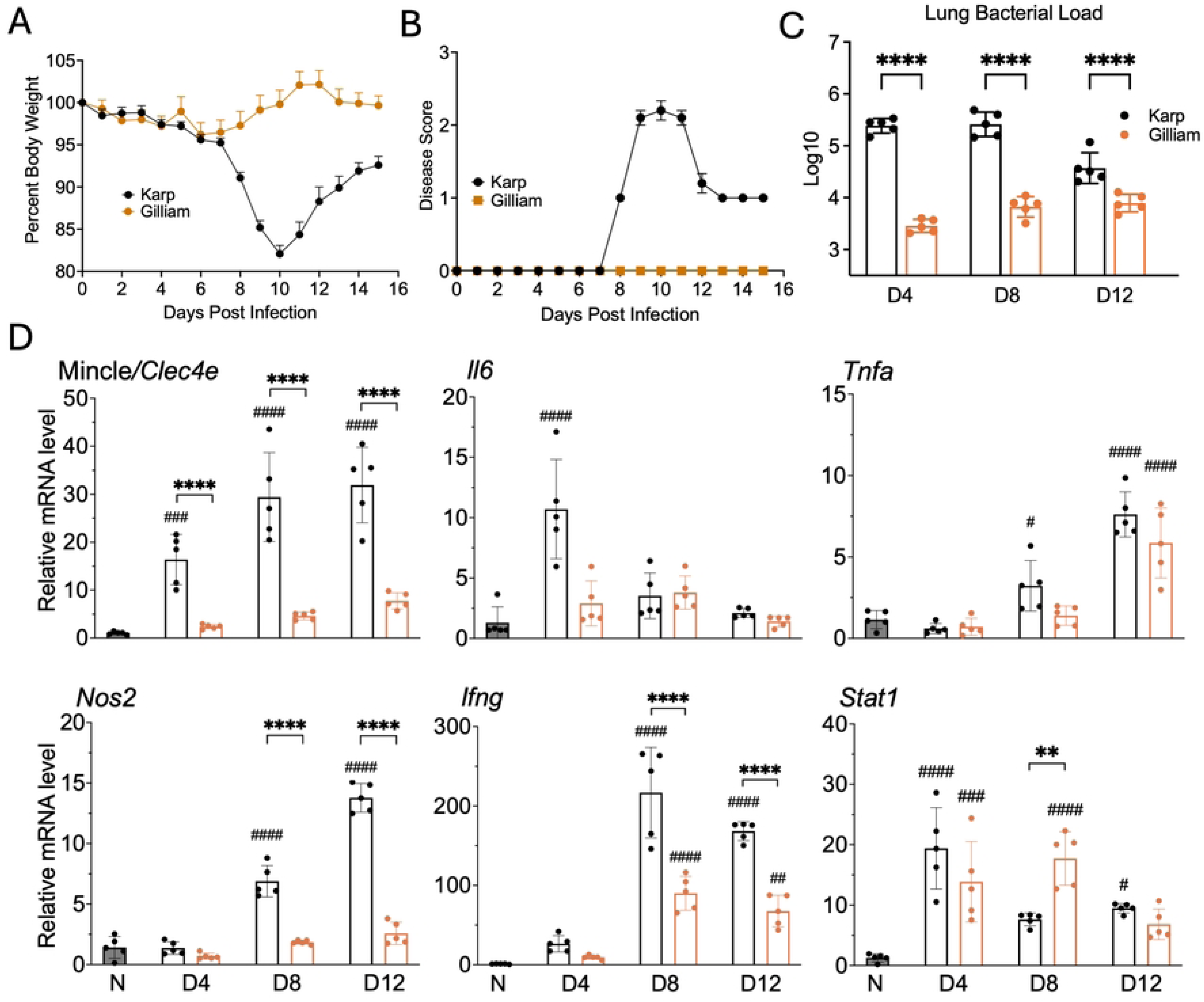
Strong upregulation of type 1 inflammation-associated genes in Karp- vs. Gilliam-infected lung tissues during infection. Mice were infected with *Ot* Karp or Gilliam strain (5.6 × 10^4^ FFU, i.v.) and monitored daily (n = 5/group) for (A) the percentage of survival rates (%) and (B) disease scores. Lung tissues collected on days 4, 8, and 12 post-infection were used for (C) qPCR analyses of bacterial loads, showing copy numbers per ng of tissue DNA on a log10 scale, and (D) qRT-PCR analyses of the indicated genes, showing relative mRNA levels normalized to those of *Gapdh* transcripts. Data are presented as mean ± SD. Statistical significance regarding the uninfected mice (N) was determined using One-way ANOVA with Dunnett’s multiple comparison test (#*p* < 0.05, ##*p* < 0.01, ###*p* < 0.001, ####*p* < 0.0001). Statistical significance was determined by comparing Karp and Gilliam at each time point using one-way ANOVA with Šídák’s multiple comparisons test (\*\**p* < 0.01, \*\*\*\**p* < 0.0001).

## Results

### Karp Infection Strongly Activates Inflammatory MΦ-related Genes in Lung Tissues

To define disease outcomes and early immune responses during infection, we infected B6 mice with a similar infectious dose of Karp or Gilliam strain (5.6 × 10^4^ FFU, i.v.). Consistent with our previous report (17), Karp-infected mice had increased disease scores and progressive body weight loss around 8 dpi (Fig. 1A-B), as well as consistently high lung bacterial burdens at 4, 8, and 12 dpi (Fig. 1C). In contrast, Gilliam infection was self-limited with no disease signs. To further test our hypothesis that such distinct disease outcomes are regulated by the innate immunity, especially MΦ-related immune responses (17), we examined mRNA levels of MΦ-associated proinflammatory markers in lung tissues via qRT-PCR. As shown in Fig. 1D, while both *Il6* and Mincle/*Clec4e* mRNA levels were significantly higher in Karp than in Gilliam infection at 4 dpi, Mincle/*Clec4e* levels continually increased in Karp group at 8 and 12 dpi; however, their levels in Gilliam-infected lungs were not significant at any time point as compared to mocks. While *Ifng* gene was updated at 8 and 12 dpi in Karp and Gilliam groups, its levels in Karp group were significantly higher, which positively correlated with *Nos2* expression levels. Again, *Nos2* levels in Gilliam-infected lungs were not significant at 8 and 12 dpi as compared to mocks. In contrast, expression profiles of *Stat1* (the JAK/STAT signaling downstream of IFNs (36)) and *Tnfa* seemed to be comparable between two *Ot* groups. These data suggested that differential activation of innate immune genes in infected lung tissues can influence the host susceptibility to Karp vs. Gilliam infection.

### Karp Infection Induces a Proinflammatory Gene Profile, While Gilliam Infection Distinctively Upregulates Cell Division and Cell Cycle-Related Genes in MΦs

Given our above findings of distinct disease outcomes and selective activation of innate responses, we wanted to examine what innate genes/pathways are selectively activated in Karp- vs. Gilliam-infected MΦs. We generated primary MΦs from mouse bone marrow, infected them with Karp or Gilliam strain (MOI 10), and analyzed mRNA profiles by bulk RNA sequencing at 1 and 3 dpi. Given the relatively slow replication rates of *Ot* bacteria (37), it was not surprising that there were no/limited major transcriptional differences among groups at 1 dpi (Fig. S1); our subsequent analyses were focused on 3 dpi. As shown in Fig. 2A, Karp infection induced 1,024 DEGs (595 upregulated, 429 downregulated), whereas Gilliam infection resulted in 2,075 DEGs (1,120 upregulated, 995 downregulated; cutoff 1.25 |log2(FC)|) when compared to the uninfected control group. Several PRRs (*Tlr2,* Dectin2*/Clec4n,* Mincle/*Clec4e, Clec5a, Nod2,* etc.) were significantly upregulated in Karp-infected MΦs; in sharp contrast, many PRPs (*Tlr8, Tlr9, Clec4a1/2/3, Cled4d,* Dectin2/*Clec4n,* etc.) were predominantly downregulated in Gilliam infection. KEGG enrichment analysis further revealed distinct responses (Fig. S2A-B). Karp infection was enriched for inflammatory and PRR-related pathways, with the “TNF signaling pathway” as the second-highest enrichment. (Fig. S2A). In contrast, Gilliam infection was unexpectedly enriched for cell cycle-related processes, with “DNA replication initiation” as the top pathway (Fig. S2B).

**Figure 2.**
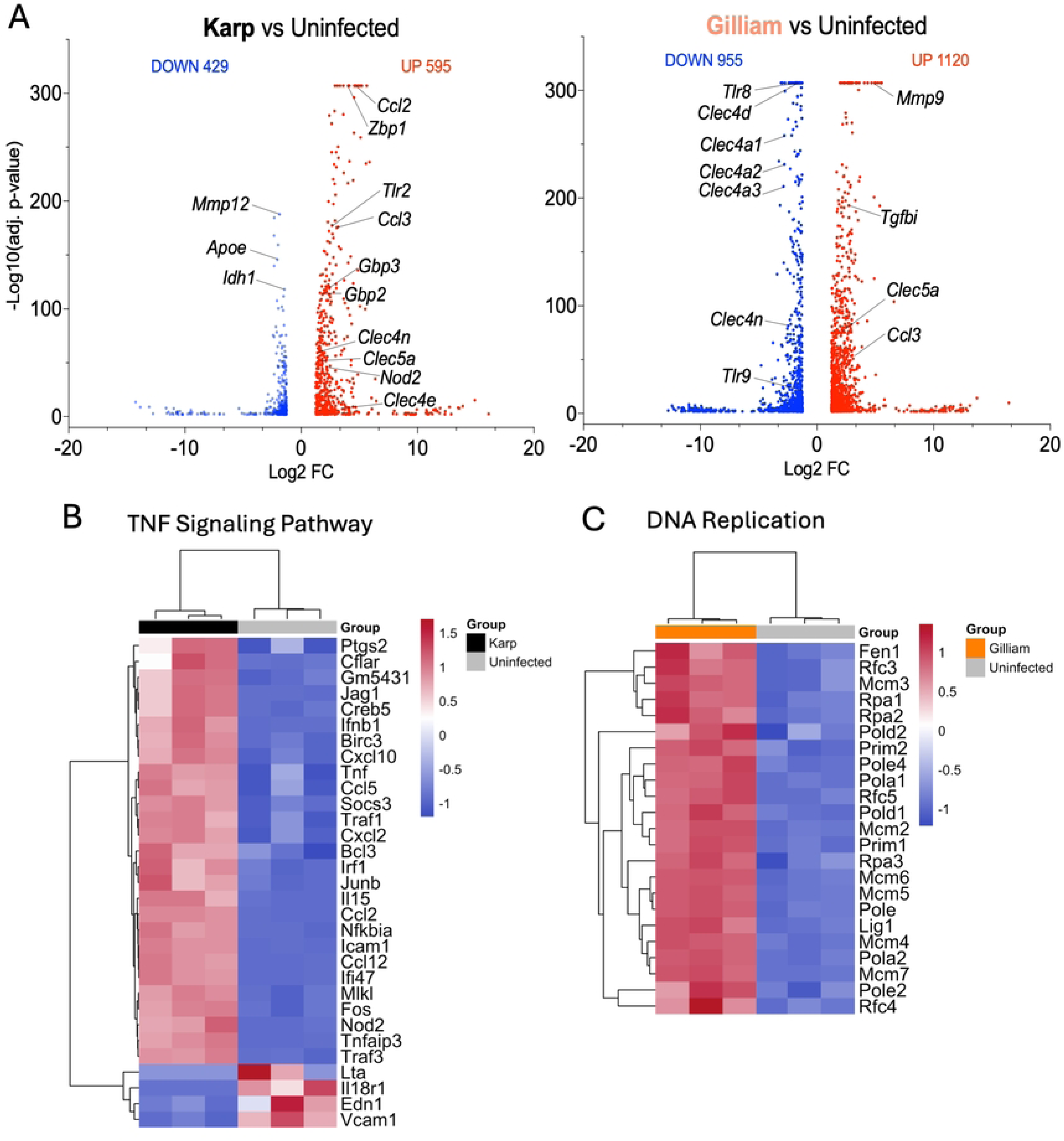
MF differential gene profiles with inflammation-skewed responses in Karp, but proliferation-associated pathways in Gilliam, infection. C57BL/6 mouse bone marrow–derived MΦs were infected with Karp or Gilliam strain (MOI 10) and harvested at 3 days post-infection (3 biological replicates per condition), alongside uninfected controls. Total RNA was subjected to high-throughput sequencing using the NovaSeq 6000 System, and differential gene expression analysis was performed using DESeq2. (A) Differential expression analysis visualized through volcano plots revealed statistically significant genes relative to uninfected samples (Karp: 429 downregulated and 595 upregulated; Gilliam: 955 downregulated and 1,120 upregulated). Genes meeting the significance threshold (|log₂FC| ≥ 1.25, -log(*p*-val) > 1.3) are shown. Heatmaps show row-scaled gene expression (Z-scores) for (B) TNF-related signaling and (C) DNA replication-related genes across infected (Karp or Gilliam) and uninfected samples. Pathways were selected based on KEGG top enrichment. Data are shown for biological triplicates from Karp-infected cells (black) and Gilliam-infected cells (orange), with hierarchical clustering applied to visualize expression patterns across conditions.

Consistent with the KEGG enrichment results, heatmap visualization of pathway-specific gene expression further highlighted the divergent transcriptional responses induced by these two strains at 3 dpi. In the TNF signalling pathway (Fig. 2B), Karp-infected MΦs exhibited robust upregulation of a broad set of inflammatory mediators, including cytokines, chemokines, and NF-κB-associated genes (*Tnf*, *Cxcl10*, *Ccl5*, *Nfkbia*, *Birc3*, *Icam1,* etc.). This coordinated induction underscored a strong proinflammatory response driven by Karp infection. Conversely, the DNA replication pathway heatmap revealed upregulation of replication machinery genes in Gilliam-infected MΦs, including components of the minichromosomal maintenance complex, DNA polymerases, and replication factors (Fig. 2C). Table 1 highlighted a side-by-side comparison between Karp and Gilliam strains for the expression of 10 PRRs, 7 signalling molecules, and 8 cytokines/chemokines (with most important markers marked in bold). Together, these data for infection vs. mock comparison illustrate that Karp infection preferentially activated inflammatory and innate immune signalling pathways, whereas Gilliam infection promoted a transcriptional profile associated with DNA replication and cell cycle progression.

**Table 1.**
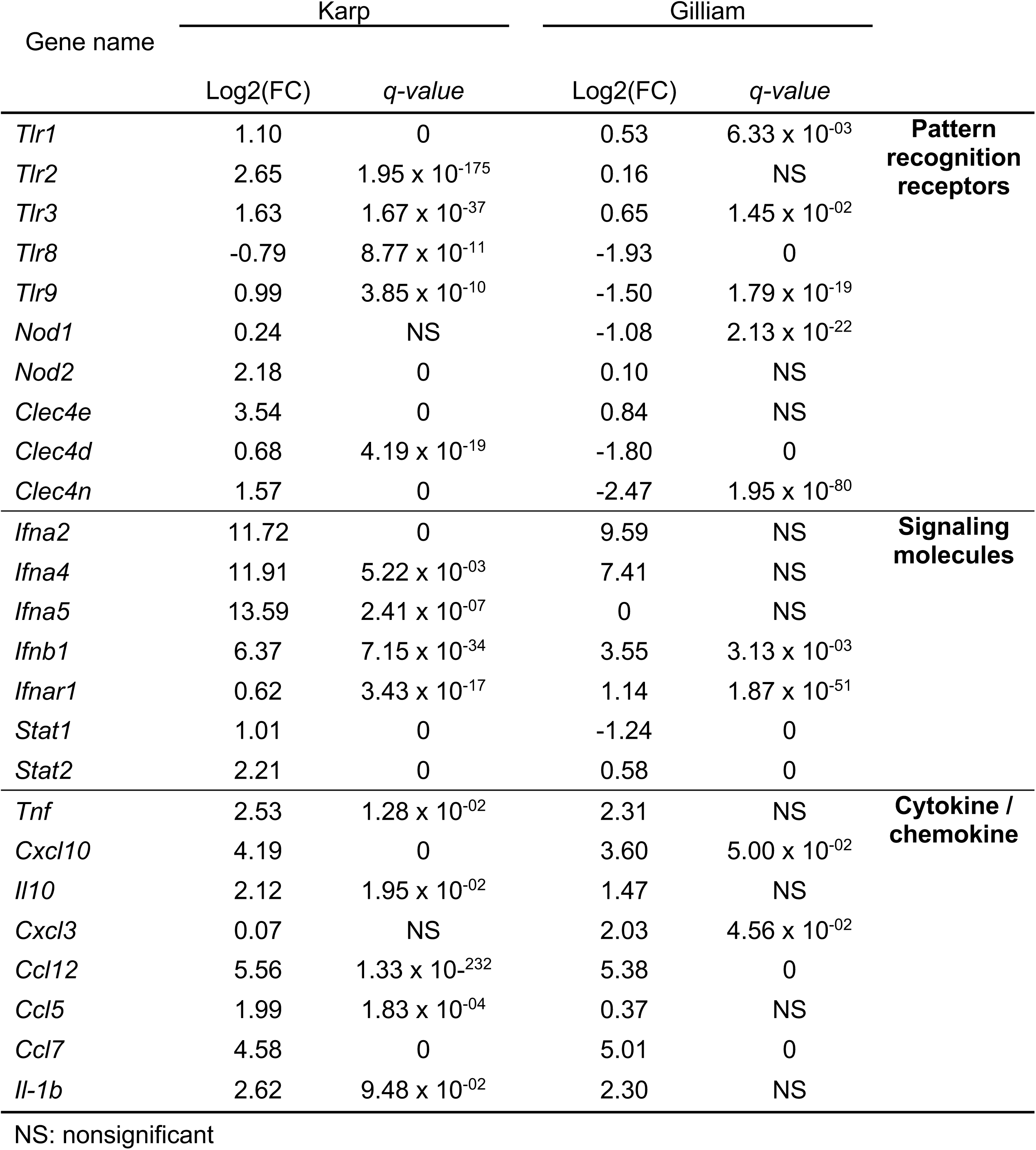
Differential immune gene expression in Karp vs. Gilliam infection.

### Strain-Specific Divergence in PRR Signaling Gene Expression Profiles in Karp- vs. Gilliam-Infected MΦs

Having demonstrated *Ot* infection-associated gene regulation in each strain, we then performed Karp-vs-Gilliam comparisons. As shown in Fig. 3A, differential expression analysis identified a total of 2,266 DEGs, including 1,237 downregulated and 1,029 upregulated genes. Notably, comparative analysis of PRR gene expression revealed that Karp infection induced higher overall expression levels than Gilliam, with NLRs, TLRs, and CLRs as the most significantly upregulated PRR-associated genes in Karp. At the pathway level, KEGG enrichment revealed a significant difference (*p*< 0.05) in NLR and TLR signaling components in Karp compared to Gilliam, including *Nod1/2*, Ripk3, interferon-stimulated genes (*Ifna5, Ifnb1,* and members in the *Oas* family), and downstream effectors (*Stat1/2, Irf7*) (Fig. 3B-C). In contrast, Gilliam showed reduced expression of these innate sensor-related modules but selectively induced calcium-associated genes (*Ppp3cc*, *Itpr3*). Consistently, TLR (*Tlr2, Tlr9*) and CLR signaling components were broadly elevated in Karp. Consistent with this observation, IPA comparing Karp and Gilliam infections demonstrated enrichment of the “PRRs in Recognition of Bacteria/Viruses” pathway in Karp compared to Gilliam. Karp infection was also associated with increased activation of immune and inflammatory pathways, whereas Gilliam infection preferentially enriched pathways related to cell division (Fig. 3D). Normalization of Karp and Gilliam comparison using uninfected data also showed upregulation of PRR-related pathways in Karp-infected cells (Fig. S3A-B).

**Figure 3.**
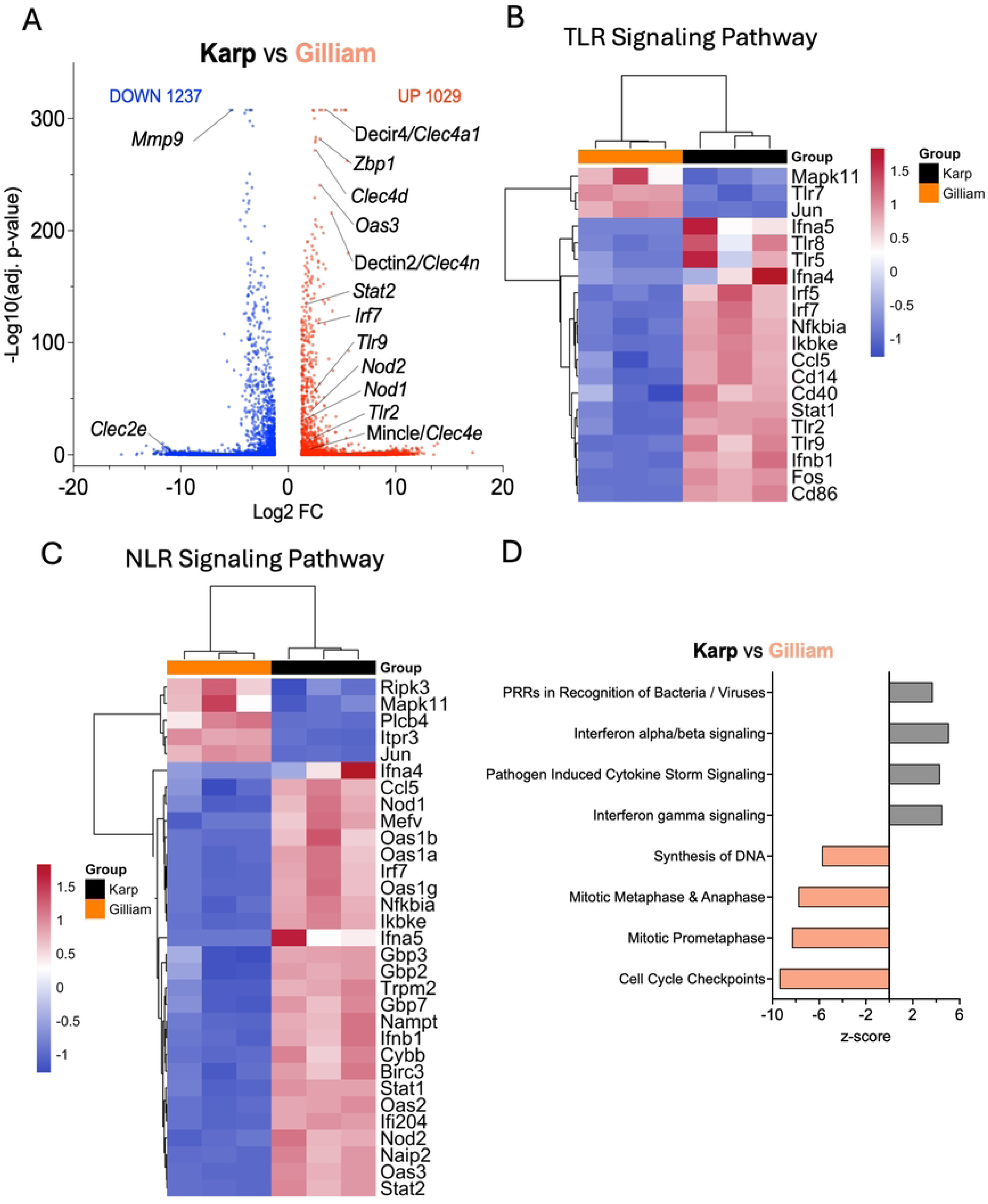
Distinct and *Ot* strain-specific gene expression programs induced in MΦ. Mouse bone marrow–derived MΦs were infected and analyzed for gene expression profiles, as in Fig. 2. (A) Volcano plot showing differentially expressed genes for Karp-vs. Gilliam-infected cells. The x-axis represents log₂ fold change (Karp vs. Gilliam), and the y-axis represents –log₁₀(adjusted *p*-value). Genes meeting the significance threshold (|log₂FC| ≥ 1.25, -log(*p*-val) > 1.3) are shown, with upregulated genes in red and downregulated genes in blue. Select differentially expressed pattern recognition receptor genes are labeled. Heatmap showing z-score–normalized expression of genes associated with (B) TLR-related and (C) NLR-related signaling pathways, selected based on KEGG enrichment and differential expression analyses. Data are shown for biological triplicates from Karp-infected cells (black) and Gilliam-infected cells (orange), with hierarchical clustering applied to visualize expression patterns across conditions. (D) Ingenuity Pathway Analysis of differentially expressed genes between Karp and Gilliam infections. The top enriched canonical pathways are shown based on –log₁₀(adjusted p-value). Activation z-scores indicate predicted pathway activity, with positive z-scores reflecting pathway activation and negative z-scores indicating inhibition, corresponding to relative enrichment in Karp or Gilliam infection, respectively.

### *Ot* Strain-Dependent Regulation of Cytokine, Chemokine, and Antibacterial Gene Expression in Infected MΦs

With the delineation of MΦ transcriptional responses to *Ot* strain-specific infection, we validated expression of selected inflammatory mediators in bone marrow-derived MΦs via qRT-PCR analyses at 1 and 3 dpi. As shown in Fig. 4A, while *Tlr2, Il1b, Gbp2,* and *Gbp3* genes were upregulated by both strains at 1 dpi, sustained upregulation of these genes was observed only in Karp-infected cells, indicating prolonged inflammatory activation. *Il12b* showed strain- and time-dependent regulation, with induction by Gilliam at 1 dpi (∼4-fold) but a stronger upregulation by Karp at 3 dpi (∼23-fold). While proinflammatory cytokines (*Il6, Tnfa, Ccl3, Cxcl2*) were not induced at 1 dpi (Fig. 4B), they were all upregulated in Karp-infected cells at 3 dpi, whereas only *Ccl3* was modestly induced by Gilliam. Notably, Karp elicited significantly higher *Cxcl2* expression compared to Gilliam (∼23-fold).

**Figure 4.**
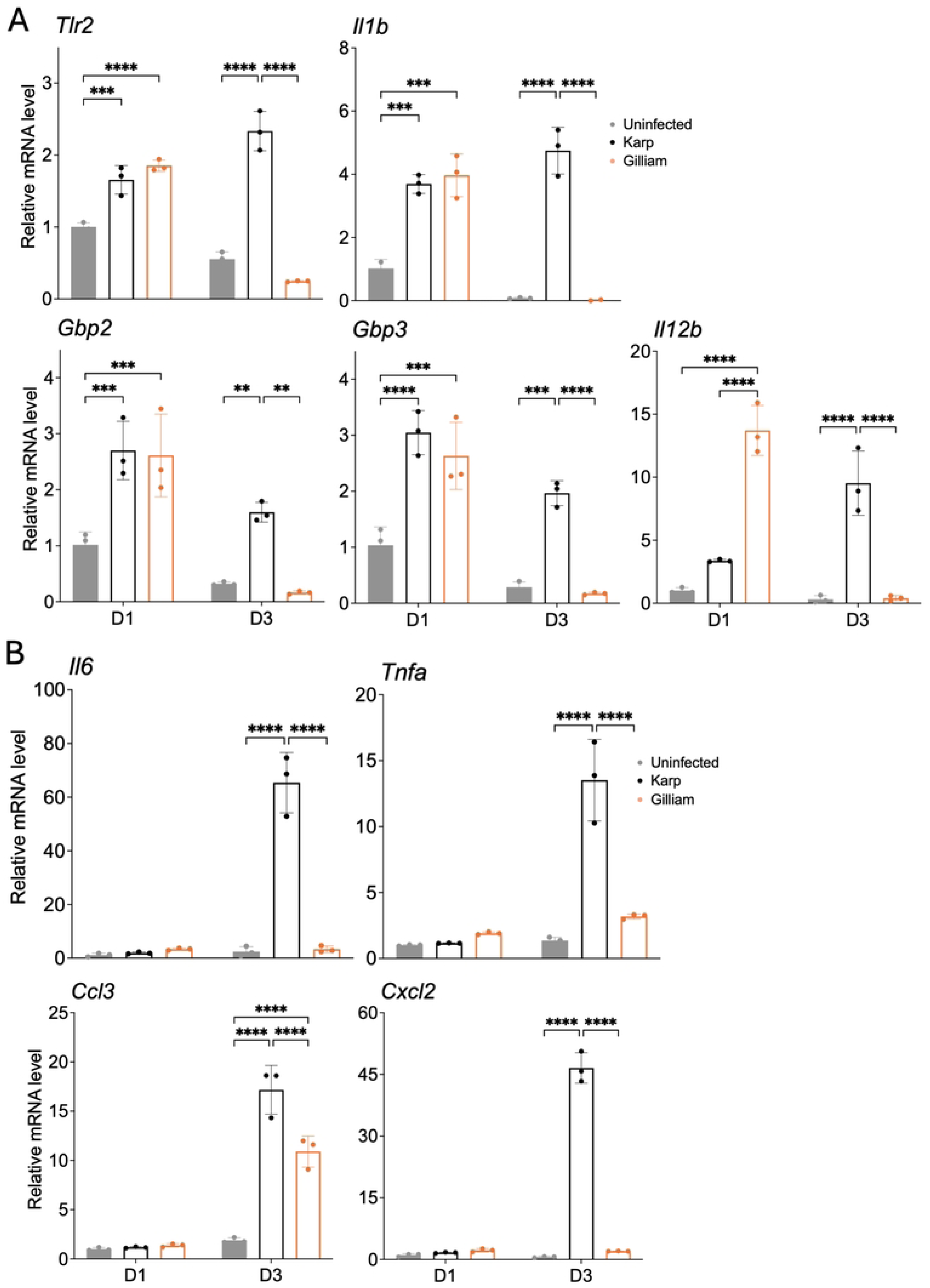
Validation of inflammatory gene expression during *Ot* Karp vs. Gilliam infection. C57BL/6 mouse bone marrow–derived MΦ were infected with *Ot* Karp or Gilliam (MOI 10) and harvested at 1 and 3 days post-infection (D1 and D3), alongside uninfected controls (3 biological replicates per condition). Total RNA was extracted for gene expression analysis by qRT-PCR. Relative mRNA levels were normalized to *Gapdh* expression. (A) Expression of *Tlr2*, *Il1b*, *Gbp2*, *Gbp3*, and *Il12b*. (B) Expression of *Il6, Tnfa, Cxcl2, and Ccl3*. Data are presented as mean ± SD of biological triplicates. Statistical significance was determined using two-way ANOVA followed by Tukey’s multiple comparisons test, with comparisons restricted within each time point (\**p* < 0.05, \*\**p* < 0.01, \*\*\**p* < 0.001, \*\*\*\**p* < 0.0001).

### M1-type MΦ Polarization Modulates Strain-specific Bacterial Replication and Upregulates CLR Signaling Gene Expression

Having demonstrated strong type 1-skewed responses in Karp infection, we next tested whether Karp and Gilliam strains react to M1-like MΦs differently. We pre-treated primary MΦs with IFN𝛾 (100 ng/ml) for 24 h, infected cells with Karp or Gilliam at different MOIs (2, 10), and harvested cells at 3 dpi (Fig. 5A). As shown in Fig. 5B, while Gilliam replicated well in untreated cells, its growth was drastically reduced in M1-like MΦs (with 94% reduction). While Karp also showed significant growth reduction in M1-like MΦs, their levels of reduction were about 60%. Importantly, qRT-PCR analyses revealed infection dose- and IFN𝛾-mediated stimulation of Mincle/*Clec4e* and Dectin2/*Clec4n* in Karp-infected, but not Gilliam-infected, cells. The expression of *Tnfa* and *Malt1* (a gene coding for a molecule involved in downstream CLR signaling) in Karp infection seemed to be consistently higher in Karp groups compared with Gilliam groups. The *Gbp2* and *Gbp3* levels were significantly higher in cells exposed to IFN𝛾 and Gilliam, especially at MOI 10 (Fig. 5C). These results suggest that M1-type polarization differentially restricted strain-specific bacterial replication, with Gilliam exhibiting greater susceptibility to IFN𝛾-mediated control than Karp. The partial resistance of Karp correlated with enhanced induction of CLR signaling components, including Mincle/*Clec4e*, Dectin2/*Clec4n*, and *Malt1*, as well as elevated *Tnfa* expression. In contrast, increased *Gbp2* and *Gbp3* expression in IFN𝛾-treated, Gilliam-infected cells suggested a more prominent role for GBP-mediated antimicrobial pathways in controlling this strain.

**Figure 5.**
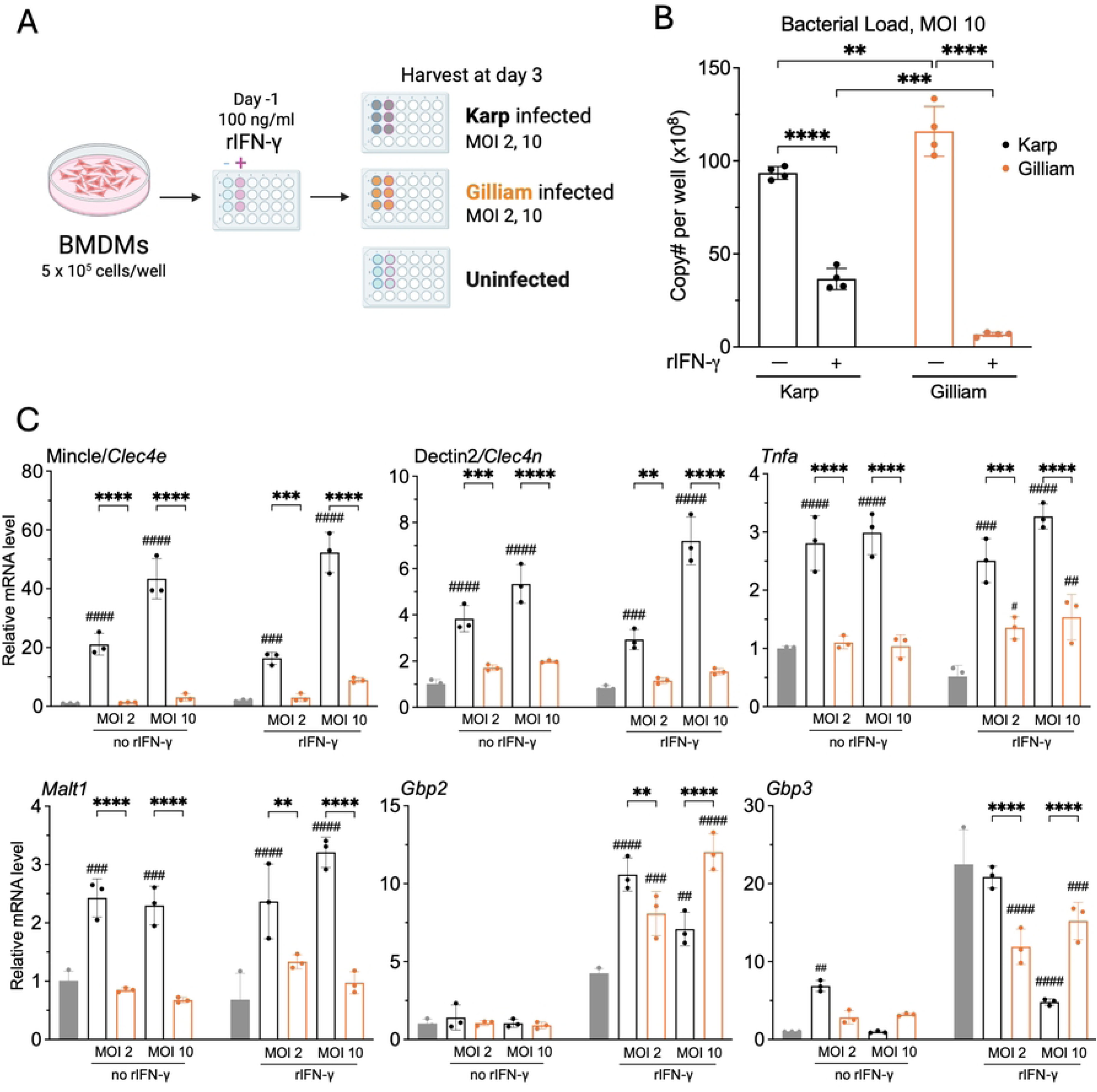
*Ot* Karp and Gilliam replication dynamics and responses in IFN-γ–primed MΦ. (A) Schematic overview of the experimental design. C57BL/6 mouse bone marrow–derived MΦs (pretreated with IFN-γ for 24 h or untreated) were infected with Karp or Gilliam (MOI 2 and 10). Samples were collected at 3-dpi. (B) Quantification of bacterial burden in Karp- and Gilliam-infected MΦ by qPCR, expressed as bacterial DNA copy number per well (×10^8^). Data are presented as mean ± SD (n = 4 biological replicates). Statistical significance was determined using one-way ANOVA followed by Tukey’s multiple comparisons test. (\**p* < 0.05, \*\**p* < 0.01, \*\*\**p* < 0.001, \*\*\*\**p* < 0.0001). (C) qRT-PCR analysis of *Clec4e, Clec4n, Tnfa, Malt1, Gbp2, and Gbp3* expression in Karp- or Gilliam-infected MΦ at 3-dpi. Relative mRNA levels were normalized to *Gapdh* and are presented as mean ± SD (n = 3 biological replicates). Statistical significance was determined using two-way ANOVA with Tukey’s multiple comparisons test. Symbols indicate comparisons between strains (*) or relative to uninfected controls (#), with comparisons restricted within the same time point (\**p* < 0.05, \*\**p* < 0.01, \*\*\**p* < 0.001, \*\*\*\**p* < 0.0001; #*p* < 0.05, ##*p* < 0.01, ###*p* < 0.001, ####*p* < 0.0001).

## Discussion

Despite being an important and emerging neglected tropical disease, scrub typhus remains understudied, particularly with respect to how strain-level variation in *Ot* shapes host innate immune responses and disease severity. Our previous reports have revealed that Karp and Gilliam strains differ markedly in pathogenicity in B6 and CD-1 mice, with Karp causing more severe disease and stronger systemic inflammatory responses than Gilliam (17, 34). In this study, we provided new evidence that these divergent disease phenotypes are associated with fundamentally distinct MΦ transcriptional programs. Specifically, Karp infection drives a coordinated proinflammatory innate immune response characterized by broad activation of PRR pathways; however, Gilliam infection induces a comparatively attenuated inflammatory phenotype accompanied by unexpected enrichment of host cell cycle and DNA replication pathways. These findings of *Ot* strain-specific innate immune programming as a central determinant of *Ot* pathogenesis provide a framework linking differential immune sensing to divergent disease outcomes.

One of our major findings from infected lung tissues and different MΦ culture conditions is that strain-specific divergence begins at the level of innate pathogen sensing. Karp infection induced strong upregulation of multiple PRR families, TLRs, NLRs, and CLRs, suggesting that this strain engages MΦ innate sensors in a broad and coordinated manner. Among these, TLR2 is particularly important because it is the only TLR experimentally shown to recognize *Ot* ligands, and its activation has been associated with increased inflammatory cytokine production and enhanced susceptibility to murine scrub typhus rather than protection (2). In our study (Figs. 2A, 4A), sustained *Tlr2* upregulation in Karp-infected MΦ correlated closely with increased expression of canonical inflammatory mediators, including *Tnf, Il1b, Il6, Ccl3,* and Cxcl2, consistent with activation of TNF/NF-κB signaling pathways. The higher expression of *Tlr9* in Karp infection was also notable and relevant, as TLR9 (a key intracellular DNA sensor) is known to recognize bacterial CpG motifs, involved in intracellular bacterial infections (ehrlichiosis, salmonellosis, etc.), modulate inflammasome activation, and influence disease severity (38–40). Although direct TLR9 recognition of *Ot* has not yet been demonstrated, it is possible that sensing bacterial nucleic acids may contribute to the heightened inflammatory phenotype observed in this strain.

In parallel with TLR activation, Karp also preferentially upregulated *Nod1* and *Nod2*, indicating enhanced cytosolic bacterial sensing. This is particularly intriguing because *Ot* has long been considered atypical among bacteria in the *Rickettsiaceae* family due to its lack of classical peptidoglycans. However, recent evidence for peptidoglycan-like structural components in *Ot* provides a plausible molecular basis for NOD receptor engagement (3). NOD1 and NOD2 recognize peptidoglycan-derived motifs and activate RIPK2-dependent NF-κB and MAPK signaling, thereby amplifying inflammatory and antimicrobial responses (41, 42). The coordinated induction of *Nod1/2*, interferon-stimulated genes, and downstream transcription factors such as *Stat1* and *Stat2* in Karp-infected MΦ suggests that NOD signaling may act synergistically with TLR pathways to intensify inflammatory MΦ activation. Similar cooperative PRR amplification has been described in other intracellular bacterial systems (43, 44), which call for future investigation into its role in exaggerated inflammation during severe Karp infection.

CLR signaling represents the third major axis of divergence between our examined strains, and their gene expression levels are most striking in the context of Karp-associated inflammation. Our data (Figs. 1D, 5C) show marked upregulation of Mincle/*Clec4e* and Dectin2/*Clec4n* in Karp infection, both in infected MΦ and mouse lung tissues. This finding aligns with previous work showing that *Ot* selectively stimulates Mincle and promotes type 1-skewed proinflammatory immune responses (21), as well as recent evidence that *Ot* can bind multiple CLRs directly (33). Mincle and Dectin-2 signal through Syk and the CARD9-BCL10-MALT1 complex to activate NF-κB and inflammatory cytokine production (45–47). In our study (Fig. 5C), increased *Malt1* expression in Karp-infected cells further supports activation of these downstream signaling pathways. Together, these findings suggest that Karp triggers convergent activation of TLR, NLR, and CLR pathways, producing a multi-layered PRR signaling network that amplifies MΦ inflammatory output.

This broad PRR activation likely explains the strong inflammatory effector profile induced by Karp infection. Compared with Gilliam, Karp elicited robust induction of proinflammatory cytokines and chemokines, including *Il6, Tnfa, Ccl3,* and *Cxcl2*, together with enrichment of the TNF signaling pathways. Such responses are likely beneficial for bacterial control early in infection, yet excessive activation can also become pathogenic. In severe scrub typhus, exaggerated type 1 inflammation has been linked to endothelial dysfunction, vascular leakage, and tissue injury (22, 48, 49). While our previous reports have shown progressive Karp replication/dissemination, pulmonary pathology, and mouse death (10, 17, 21), this study provides transcriptomic evidence that helps understand how Karp induces innate sensing and inflammatory pathways, driving both antibacterial defense and collateral host damage. Thus, Karp appears to occupy a pathogenic niche, in which stronger innate recognition enhances immune activation with unsuccessful bacterial clearance (Fig. 1C) and increased immunopathology.

In sharp contrast, Gilliam followed a markedly different infection outcome and immunological trajectory, even though the genome-wide comparisons of multiple *Ot* strains (including Karp and Gilliam) did not identify any gene clusters responsible for differential infection outcomes (16, 50), pointing to the role of the immune response induced by each strain. We found herein that, rather than strongly activating PRR networks, Gilliam either suppressed or failed to induce many PRR-associated genes, including several TLR-, NLR-, and CLR-related modules. This attenuated innate sensing profile likely explains the weaker inflammatory phenotype observed in Gilliam-infected lung tissues (Fig. 1D) and MΦ cultures (Figs. 2A, 4). Importantly, Gilliam’s reduced inflammatory signature does not reflect passive inactivity, as it induced a higher number of differentially expressed genes than Karp. Gilliam-induced genes/enriched pathways, however, were not focused on inflammation, but on DNA replication, S-phase progression, and cell cycle regulation. This suggests that Gilliam actively redirects host transcriptional machinery away from inflammatory activation and toward alternative cellular programs that may favor host cell survival or repair.

In contrast to Karp-induced CLR pathways, Gilliam preferentially downregulated several dendritic cell immunoreceptors (Dcirs), including *Clec4a1, Clec4a2,* and *Clec4a3*, whereas Karp maintained these near baseline levels. Dcirs transmit inhibitory ITIM-mediated signals that modulate inflammatory activation (55), and their altered expression may reshape MΦ signaling thresholds during infection. Because the Dcir family members have been implicated in detrimental immune regulation during bacterial infections such as tuberculosis (51), their suppression in Gilliam infection may reflect selective remodeling of inhibitory circuits that help balance bacterial persistence with reduced inflammatory injury. Whether this contributes directly to the milder phenotype of Gilliam remains an important question for future study.

Our functional experiments with IFN𝛾-polarized MΦ further support the concept that Karp and Gilliam employ distinct survival strategies (Fig. 5). Although both strains exhibited reduced replication in M1-polarized MΦ, Gilliam was substantially more susceptible to IFN𝛾-mediated restriction, whereas Karp retained partial resistance. This suggests that once MΦ become fully activated, Gilliam lacks effective countermeasures against antimicrobial killing. By contrast, Karp may possess intrinsic mechanisms that confer relative resistance to activated MΦ defenses. One possible explanation is differential evasion of GBP-mediated antimicrobial pathways, analogous to bacterial strategies described in *Shigella flexneri*, where pathogen-derived ubiquitin ligases neutralize GBP targeting (52, 53). Whether Karp possesses analogous mechanisms remains unknown, but this represents an important opportunity for further investigation.

In summary, this study indicates that Karp drives a highly inflammatory PRR-dominant program involving coordinated activation of TLR, NLR, and CLR pathways, resulting in strong antibacterial signaling but also greater host tissue injury. In contrast, Gilliam dampens inflammatory sensing but rewires host-cell transcription toward cell-cycle regulation, likely minimizing immunopathology and promoting host cell survival. Our data collectively revealed that Karp and Gilliam represent two fundamentally distinct immunopathogenic strategies within *Ot* infection in MΦs. These results provide a unified mechanistic explanation for strain-dependent disease heterogeneity in scrub typhus, suggesting innate immune sensing pathways as promising targets for biomarkers in severe disease outcomes.

## Figure Legends

**Supplementary Figure 1.** Principal component analysis of uninfected, Karp- and Gilliam-infected MΦs based on RNA sequencing data. PCA was performed on normalized log2 RNA-seq counts from 3 biological replicates per group. Each point represents a single sample, colored and labeled according to group identity (e.g., Karp, Gilliam, uninfected). Samples from the same group cluster together, while samples from different groups separate along PC1 and PC2, indicating group-specific transcriptional differences.

**Supplementary Figure 2.** (A, B) KEGG enrichment analysis of regulated genes in Karp or Gilliam vs. Uninfected comparisons (n = 3). (C, D) Heatmap showing z-score–normalized expression of genes associated with NF-κB signaling pathway, P53 signaling pathways, TNF signaling pathway, and JAK-STAT Signaling Pathway selected based on KEGG enrichment and differential expression analyses. Data are shown for biological triplicates from infected (gray), Karp-infected cells (black), or Gilliam-infected cells (orange), with hierarchical clustering applied to visualize expression patterns across conditions.

**Supplementary Figure 3.** Comparison of commonly downregulated & upregulated canonical pathways identified by Ingenuity Pathway Analysis in Karp- versus Gilliam-infected MΦ relative to uninfected controls. Axes represent –log_10_(Benjamini–Hochberg adjusted p-values), enabling direct comparison of pathway enrichment significance between the two infection conditions.

